# Detection and genetic characterization of pigeon gammacoronavirus

**DOI:** 10.1101/2025.05.22.655467

**Authors:** Hiroko Kobayashi, Mai Kishimoto, Sakiho Imai, Yasuko Orba, Hirofumi Sawa, Masayuki Horie

## Abstract

Most members of the genus *Gammacoronavirus* infect avian hosts, but far fewer viruses have been characterized than in the other coronavirus genera, leaving their diversity largely unclear. Pigeon gammacoronaviruses were previously detected by consensus PCR, but only partial sequences were determined. In this study, we comprehensively analyzed public RNA-seq datasets and reconstructed two nearly complete genomes of pigeon gammacoronaviruses. Molecular evolutionary analyses showed pigeon gammacoronaviruses belong to the subgenus *Igacovirus*, and their pairwise distances to the members of this subgenus meet the International Committee on Taxonomy of Viruses (ICTV) species demarcation criteria, supporting their designation as a novel species in this subgenus. Notably, although nucleotide sequences are highly conserved gammacoronaviruses, their 3′-genomic region exhibited differences in gene organization among pigeon coronavirus variants. These findings expand our knowledge of the diversity of gammacoronaviruses.

## Introduction

The diversity of viruses has not yet been fully elucidated. To strengthen our preparedness for infectious diseases and to understand the evolution of viruses, it is essential to continue searching for viruses from a variety of samples. Various methods have been used to detect novel viruses. Consensus PCR employs primers that target conserved regions, enabling inexpensive, highly sensitive detection of related viruses without cell or animal culture. Consequently, it has been widely adopted and has yielded valuable insights into viral diversity [1]. However, in some cases, longer genome sequences for such viruses have not been determined. Because some viruses cause recombination, it is necessary to have longer sequences to understand the evolution of viruses. Furthermore, in many viruses, nearly complete genome sequences are required for virus classification. Thus, it is important to determine nearly complete genome sequences of viruses for which only partial sequences have been identified.

Coronaviruses are single-stranded RNA viruses belonging to the subfamily *Orthocoronavirinae* of the family *Coronaviridae* of the order *Nidovirales*. Coronaviruses infect various animals, causing respiratory, enteric, hepatic and neurological diseases [2]. Coronaviruses are further divided into four genera, *Alphacoronavirus, Betacoronavirus, Gammacoronaviurs*, and *Deltacoronavirus* [3]. Among them, the genus *Gammacoronavirus* contains viruses mainly infecting avian species except for cegacoviruses in some marine mammals [4,5], which can be further divided into three subgenera, *Igacovirus, Cegacovirus* and *Brangacovirus*. Some gammacoronaviruses are known to infect poultries and cause respiratory disease and decrease in egg production [6]. They are also detected in a wide range of wild birds, such as rock pigeons (*Columba livia*) [7,8]. A previous study first detected gammacoronavirus from tracheal and cloacal swabs, liver, and spleen of rock pigeons in Norway, which were apparently healthy with no pathological lesions were found at necropsy [9]. Further, pigeon gammacoronaviruses have been found from feral and domestic pigeons in Finland [10], Poland [11,12], and China [13,14]. However, because the viruses were detected using consensus PCR, only partial sequences are available in the database.

In this study, we aimed to obtain a nearly complete genome sequence of pigeon gammacoronavirus. To do so, we analyzed publicly available RNA-seq data and identified two nearly complete genomes of pigeon gammacoronavirus. Comparative genome analysis revealed that the pigeon gammacoronavirus has unique genome structure among the members of subgenus *Igacovirus*. Further, pigeon gammacoronavirus meets the International Committee on Taxonomy of Viruses (ICTV) species demarcation criteria, suggesting that the virus can be a novel species within the subgenus *Igacovirus*. Thus, this study provides a novel insight of the diversity and evolution of avian gammacoronavirus and highlights the importance of further analysis of viruses for which only partial sequences have been determined.

## Materials and Methods

### Detection of pigeon coronavirus-related contigs in public RNA-seq data

To identify RNA-seq datasets containing pigeon coronavirus-derived sequences, 1,224 RNA-seq datasets of *C. livia* (Table S1) were mapped to the sequence of pigeon coronavirus isolate 03/653 (accession number: AJ871022) [9] using Magic-BLAST 1.7.1 [15] with the default setting except for the option “-word_size 16”.

To reconstruct genomes sequences of pigeon gammacoronaviruses, RNA-seq datasets with more than 1,000 mapped reads against the pigeon coronavirus sequence (in total 94 datasets; Table S2) were downloaded from NCBI SRA [16] and preprocessed by fastp 0.20.1 using the “-l 35 -x” options [17]. The preprocessed reads were then mapped to the reference genome of *C. livia* (GenBank assembly accession: GCA_028654425.1) by HISAT 2.1.0 [18]. The unmapped reads were extracted using SAMtools 1.15.1 [19] and assembled by rnaviralSPAdes 3.15.5 with the default setting [20]. The obtained contigs more than 10,000 nucleotides were extracted using SeqKit [21]. To extract coronavirus-like contigs, sequence similarity search was performed against coronavirus sequences in the NCBI nr database (taxid 11118; *Coronaviridae*) by BLASTx 2.9.0 [22] using the extracted contigs as queries with the following options: -evalue 1e-4, -max_target_seqs 10. The BLAST results were manually analyzed.

To further detect RNA-seq datasets containing pigeon coronavirus-derived sequences using nearly complete sequence of pigeon gammacoronavirus (accession number BR002451), Magic-BLAST was performed using the 1,224 RNA-seq datasets against a pigeon coronavirus contig sequence as describe above. Then, 256 RNA-seq datasets containing pigeon gammacoronavirus reads (Table S3) were subjected to *de novo* assembly as described above. Some contigs were further assembled using Geneious assembler in Geneious Prime v2024.0.5 (https://www.geneious.com). Note that some RNA-seq datasets, which were obviously obtained from the same samples based on the metadata, were merged before assembly (Table S4). Among the obtained contigs, those encoding the full-length S and/or N gene were extracted by BLASTn and manual analysis, which were used for downstream analyses.

To validate the obtained pigeon gammacoronavirus contigs, the original unmapped reads were mapped back to the contigs, and the coverage depths were calculated using SAMtools. Sequences covered by less than three reads were removed.

All the contigs used in this study were deposited in DDBJ under accession numbers BR002420- BR002451 (Table S4).

### Annotation of pigeon gammacoronavirus contigs

Open reading frames (ORFs) more than or equal to 150 nucleotides were detected by the “Find ORFs” function in Geneious Prime. To annotate the ORFs, BLASTx was performed against the NCBI nr database using the nucleotide sequences of detected ORFs as queries. Three ORFs, which did not show any BLAST hits, were annotated as ORF4c, ORFY-1 or ORFY-2. The coding region of ORF1ab was deduced considering -1 programmed ribosomal frameshift on the slippery sequence (TTTAAAC) [23].

### Phylogenetic analyses

Representative gammacoronavirus sequences, which are listed in the ICTV chapter for the family *Cornaviridae* [3], were manually downloaded and their ORF1b sequences were extracted. Amino acid sequences of ORF1b were aligned using MAFFT v7.490 [24] with the L-INS-i algorithm and phylogenetic relationship was inferred by Maximum likelihood method using RAxML-NG 1.1.0 [25]. The best-fit substitution model, WAG+G4+F, was selected using ModelTest-NG. The reliability of trees was assessed by 1,000-bootstrap resampling using the transfer bootstrap expectation method.

To infer the evolutionary relationship among pigeon gammacoronavirus variants, the N gene sequences were aligned using the “Translation Align” function in Geneious Prime with the L-INS-i algorithm implemented in MAFFT. Phylogenetic trees were inferred as described above except that the substitution model TIM3+G4 was used in this analysis.

The resulting trees were visualized using MEGA7 [26].

### Comparison of amino acid sequences for species demarcation

The 3CLpro, nidovirus RdRp-associated nucleotidyltransferase (NiRAN), RdRP, and HEL1 domains of each virus was detected by InterPro search [27]. The sequences of the detected domains were extracted and concatenated. The concatenated sequences were aligned by MAFFT with the E-INS-i algorithm, and then the amino acid identities were calculated using Geneious Prime.

## Results

### Determination of nearly complete genome sequences of pigeon gammacoronaviruses using publicly available sequencing data

As described above, only some partial sequences are available for pigeon gammacoronavirus. To identify nearly complete genome sequences of pigeon gammacoronavirus, we first identified pigeon gammacoronavirus-positive RNA-seq datasets in the public database. We mapped 1,224 RNA-seq datasets of *C. livia* to the partial genome sequence of pigeon gammacoronavirus isolate 03/653, which was reported in 2005 [3], and counted the numbers of mapped reads. We found that 94 RNA-seq datasets (Table S2) contained more than 1,000 reads mappable to the partial pigeon gammacoronavirus sequence. We then assembled the RNA-seq reads and identified a contig of pigeon gammacoronavirus with nearly complete genome from SRR21669565, which shows 98.28% nucleotide identity to the previously described partial sequence (KX588632.1).

To further identify pigeon gammacoronavirus sequences, we performed a mapping-based analysis using the obtained nearly complete genome sequence. We mapped the 1,224 public RNA-seq datasets obtained from *C. livia* to the contig and found that 276 of the 1,224 SRA datasets contained sequence reads derived from pigeon gammacoronavirus (Table S3). To obtain additional pigeon gammacoronavirus contigs, we assembled the RNA-seq reads. We obtained another contig of nearly complete genome from SRR15831023. We also obtained a total of 30 partial contigs containing the N gene, the S gene, or both.

### Pigeon gammacoronavirus has a unique genome organization

We performed a detailed analysis on two nearly full-length pigeon gammacoronavirus contigs (BR002449 and BR002451 obtained from SRR15831023 and SRR21669565, respectively) (Fig. 1). The contigs are 27,804 (BR002449) and 27,792 (BR002451) nucleotides in length, respectively. The genome structures are almost identical between the two viruses, except for the presence or absence of ORF4b. ORF1ab is present in the upstream region, along with the putative slippery sequence TTTAAAC. The downstream region contains the S, E, M, and N genes. Additionally, ORF5a and ORF5b, which are conserved among gammacoronaviruses, are present between the M and N genes. Notably, there are two unique ORFs, tentatively named ORFY1 and ORFY2, that show no sequence similarity to other coronavirus genes. In contrast, these pigeon gammacoronaviruses lack ORF3a and ORF3b, which are conserved in other gammacoronaviruses.

**Figure 1.**
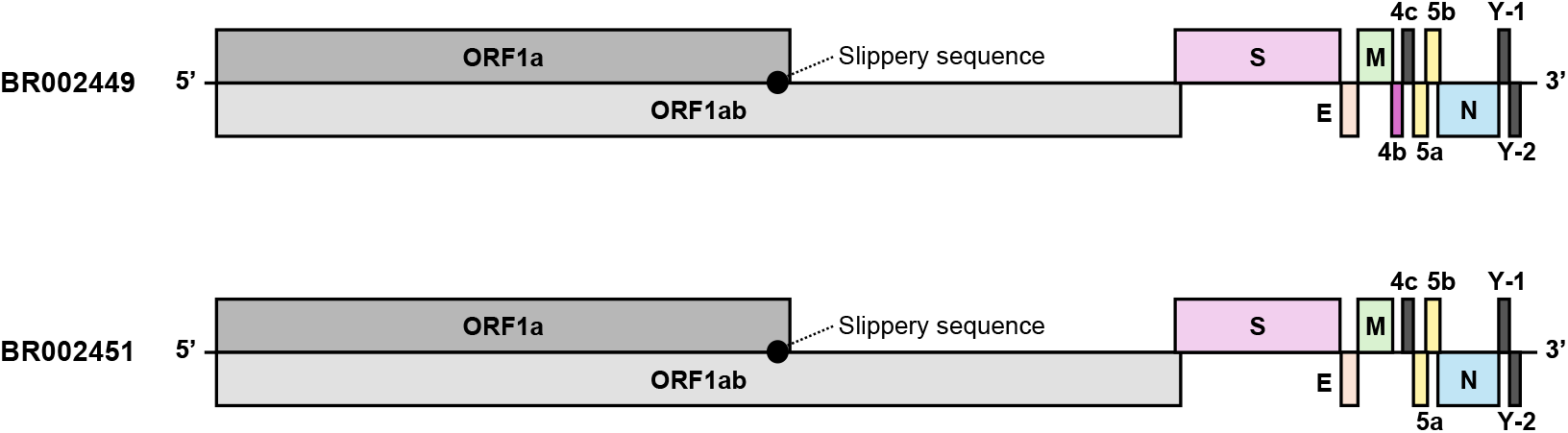
Genome structure of pigeon gammacoronaviruses. Schematic representations of genome structures of pigeon gammacoronaviruses (BR002449, BR002451) are shown. Each ORF is shown with a box. Black boxes (ORF4c, ORFY-1, and ORFY-2) indicate ORFs do not show homology to the known coronavirus genes. Black circles indicate the slippery sequences conserved among coronaviruses.

### Characterization of pigeon gammacoronaviruses

To characterize pigeon gammacoronaviruses, we analyzed the metadata of the pigeon gammacoronavirus-positive RNA-seq datasets (Table S3), revealing that these datasets originated from India and China. Furthermore, they are derived from a wide range of tissues, including the ileum, bursa of Fabricius, iris and subcutaneous fat. Notably, the virus was rarely detected in respiratory organs, such as lung (see Discussion).

### Phylogenetic analyses

To understand the evolutionary relationship between pigeon and other gammacoronaviruses, we have performed phylogenetic analyses. We first inferred the phylogenetic relationship using the ORF1b sequences of representative gammacoronaviruses (Fig. 2A). The trees showed that pigeon gammacoronaviruses form a cluster with viruses belonging to the subgenus *Igacovirus*.

**Figure 2.**
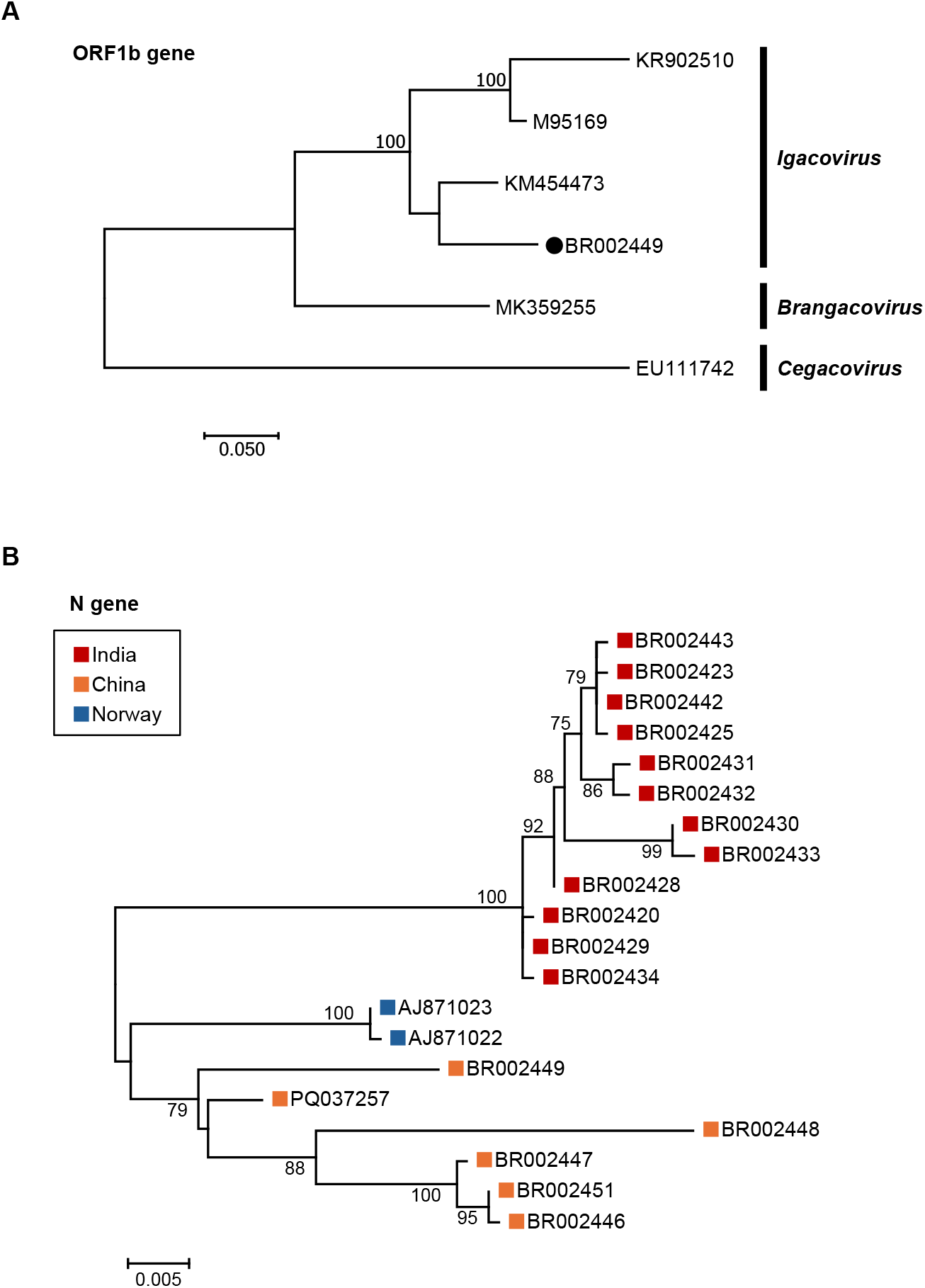
Phylogenetic relationship. **(A)** Phylogenetic tree of gammacoronaviruses constructed with the maximum-likelihood method using deduced amino-acid sequences of ORF1b proteins from representative gammacoronaviruses. **(B)** Phylogenetic relationships among pigeon gammacoronaviruses inferred with the maximum-likelihood method using nucleotide sequences of the N gene from pigeon gammacoronaviruses. Scale bars indicate the number of amino acid substitutions per site. Bootstrap values below 70 are omitted.

To further understand the diversity among pigeon gammacoronavirus variants, we performed a phylogenetic analysis based on the N gene sequences including those of previously published viruses. The tree showed that the variants are roughly divided into three clades according to the geological distribution (Fig. 2B).

### Amino acid sequence comparison for species demarcation

For species demarcation, the concatenated amino acid sequences of 3CLpro, NiRAN, RdRP, ZBD, and HEL1 domains were aligned, and sequence identities were calculated. Among the viruses compared, duck coronavirus showed the highest sequence identity (91.1%) to pigeon gammacoronavirus (Table 1).

**Table 1.**
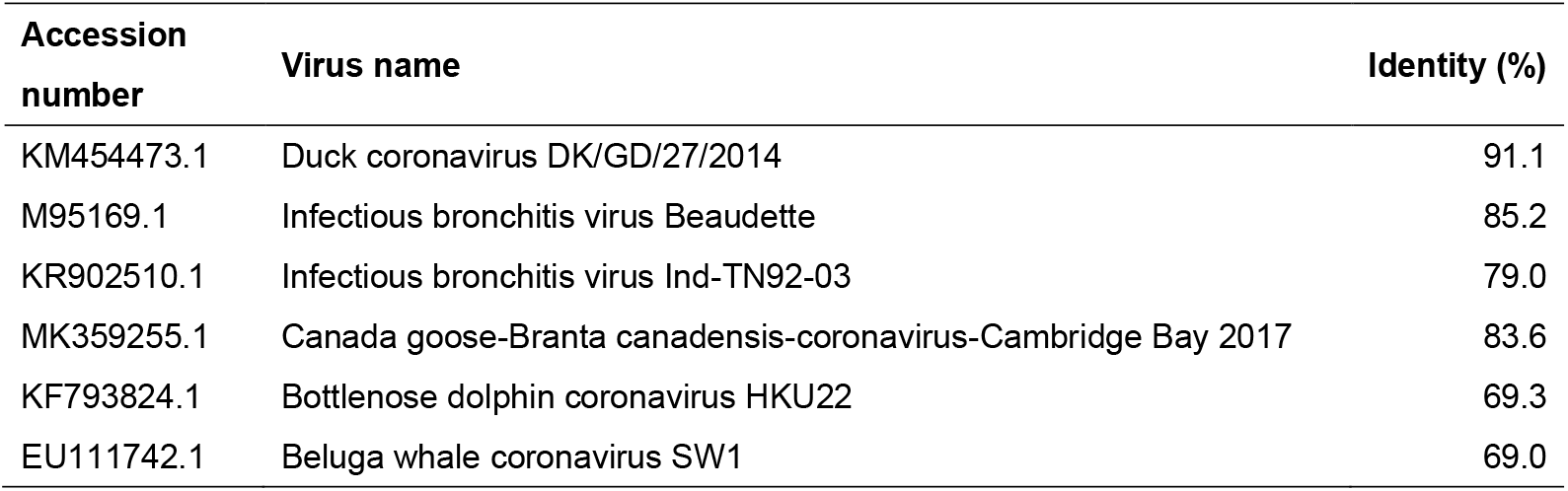
Amino acid sequence (3CLpro, NiRAN, RdRP, ZBD and HEL1 domains) identities to pigeon gammacoronavirus.

## Discussion

Although partial sequences of pigeon gammacoronaviruses have been reported thus far, research have been limited to epidemiological surveys, and the molecular characteristics of pigeon gammacoronaviruses remain largely unknown. In this study, we identified nearly complete genome sequences of pigeon gammacoronaviruses using publicly available RNA-seq datasets. The virus meet the ICTV species demarcation criteria (more than 7.5% difference in 3CLpro, NiRAN, RdRP, ZBD and HEL1 domains) [9–14], and therefore can be a novel species in the genus *Gammacoronavirus*.

Moreover, the viruses have unique genome organizations within the genus *Gammacoronavirus*. Therefore, our study provides novel insights into the diversity and evolution of gammacoronaviruses and highlights the importance of further analysis of viruses for which only partial sequences have been determined.

This study reconfirmed that pigeon gammacoronavirus is widely distributed worldwide. Previous studies have detected pigeon gammacoronaviruses in Finland, Poland, and China [9–14], suggesting an extensive geographic distribution. In addition, we detected this virus in pigeons from India in this study. These findings indicate that pigeon gammacoronavirus has an extensive geographic distribution, spanning at least from Europe to East Asia. Further studies would reveal an even broader distribution range of this virus.

Our results suggest that pigeon gammacoronavirus can infect various tissues throughout the body and may exhibit greater tropism for the intestinal tract than for the lungs. Our mapping analysis detected pigeon gammacoronavirus-derived reads in multiple tissues (Table S3). The detection of substantial numbers of reads across a wide range of organs suggests that this virus possesses broad tissue tropism and/or may cause systemic infection. Interestingly, the datasets from BioProject PRJEB41063 contain data obtained from the brain, lung, and ileum of multiple individuals. In all these set samples, the proportion of mapped reads was considerably higher in the ileum than in the lungs, and no reads were detected in the brain samples. However, it cannot be ruled out that experimental conditions or the stage of infection may have influenced these findings. Further detailed investigations focusing specifically on tissue tropism are warranted.

Together, our study identified the nearly complete genome sequences of two pigeon gammacoronaviruses and broadened our understanding of gammacoronavirus diversity. In addition, this study highlights the importance of determining nearly complete sequences of viruses for which only partial sequences are known. Further studies, such as isolation of the virus and the accumulation of genome sequence information, would facilitate understanding the diversity of gammacoronaviruses.

## Acknowledgements

The super-computing resources was provided by the Human Genome Center, the Institute of Medical Science, the University of Tokyo. This study was supported by JSPS KAKENHI grant numbers JP23K20902 (MH), JP22K19234 (MH), JP21H01199 (MH), 24K18455 (MK) and JP22J21842 (HK), the Japan Agency for Medical Research and Development (AMED) under Grants JP223fa627005 (HS), the 2024 Osaka Metropolitan University (OMU) Strategic Research Promotion Project (Young Researcher) (MK), and the World-leading Innovative and Smart Education (WISE) Program (1801) from the Ministry of Education, Culture, Sports, Science, and Technology, Japan.

